# Information-theory analysis of mouse string-pulling agrees with Fitts’s Law: Increasing task difficulty engages multiple sensorimotor modalities in a dual oscillator behavior

**DOI:** 10.1101/2023.07.13.548852

**Authors:** Pardeepak S. Sandhu, Behroo Mirza Agha, Samsoon Inayat, Surjeet Singh, Hardeep S. Ryait, Majid H. Mohajerani, Ian Q. Whishaw

## Abstract

Mouse string pulling, in which a mouse reels in a string with hand-over-hand movements, can provide insights into skilled motor behavior, neurological status, and cognitive function. The task involves two oscillatory movements connected by the string. The snout tracks the pendulum movement of the string produced by hand-over-hand pulls and so guides the hands to grasp the string. The present study examines the allocation of time required to pull strings of varying diameter. Movement is also described with end-point measures, string-pulling topography with 2D markerless pose estimates based on transfer learning with deep neural networks, and Mat-lab image-segmentation and heuristic algorithms for object tracking. With reduced string diameter, mice took longer to pull 60cm long strings. They also made more pulling cycles, misses, and mouth engagements, and displayed changes in the amplitude and frequency of pull cycles. The time measures agree with Fitts’s law in showing that increased task difficulty slows behavior and engages multiple compensatory sensorimotor modalities. The analysis reveals that time is a valuable resource in skilled motor behavior and information-theory can serve as a measure of its effective use.

## Introduction

A remarkable number of string-pulling tasks have been administer to a remarkable number of animal species, including, primates, birds and bumble bees [Alem, 2016; Jacobs ans Osvath, 2015; Riemer et al., 2014; Riesen et al, 1953; Vince, 1961]. No doubt because of their role as a laboratory species, rodents have also been featured in string-pulling tasks. The first report was a story in Life Magazine of the rat Pliny, trained by BF Skinner to pull a chain to release a marble. Pliny was reported to have pulled the chain using its teeth but photographs show that it also used its hands (Iversen, 1992). Modifications of the string-pulling task have had rats solving configural tasks involving combinations of string diameter and odor to provide insights into the sensory discrimination and learning (Tomie and Whishaw, 1990; Whishaw and Tomie, 1991; Whishaw et al, 1992). Such problem-solving ability related to string-pulling suggests that the task can be described as a rodent proto-tool task; that is, a task in which a string is used as a tool to achieve a food attainment goal. A recent series of studies with mice, rats and humans have had subjects pull down a string that is looped from an overhead track (Blackwell, 2018a; Inayat et al, 2020; Singh et al, 2019). This version of the task has advantages in that the subjects stand upright in one location, their front and hands are exposed for optimal movement tracking, allowing the rhythm and timing of hand-over-hand pulls to be subject to analyses. An additional advantage of the standing string-pulling task is that similar iterations given to mice, rats and humans provides a cross species analogue for addressing neuroscience problems (Darevsky et al, 2023; Blackwell et al, 2018b; 2021; 2022; Singh et al, 2019; Vishwanath, 2021).

Mice string-pulling is a product of the oscillatory movements of the snout and the hands connected by the string. The hands produce back-and-forth string movements that are tracked by the snout, and snout motion in turn guides subsequent hand placement on the string (Blackwell et al, 2018a). Because the string is integral in signalling these oscillatory movements, one approach to understanding their timing is to vary the information provided by the string. For example, target size generally poses a challenge in the selection of motor strategy. When feeding, most species of nonprimate terrestrial vertebrates obtain food items of small size by using the mouth to pick them up (Iwaniuk et al., 2000; Sustaita et al, 2013; Whishaw and Karl, 2019; Peckre et al 2019) whereas anthropoid primates resort to precision hand grasps to achieve the same end (Macfarlane and Graziano 2009; Napier, 1980; Hirsche et al, 2022). Accordingly, the hypothesis underlying the present study is that string size should be related to efficacy in hand use advancing the string as measured by time. Measures of time have been used to identify decision-related process associated with movement onset and task-completion processes (for example, see Guillemin, 1996; Schmidt and Lee, 2020). Furthermore, Fitts (1954) has proposed a metric for task performance centered on time, in which the average rate of information generated by a series of movements is the average information per movement divided by the time per movement. Fitts’s Law is primarily applied in the field of human motor control and there is limited research specifically linking Fitts’s Law to animal behavior. We hypothesised that based on Fitts’s metric of task difficulty in a formative task in which the center of a target is considered information and its width noise, an analogous task presented to mice is the diameter of the string for which they reached.

The dependent measures were time taken to pull in a string and qualitative and quantitative movements of advancing the strings of varying diameter. Mice that were well trained in string-pulling were presented with strings of five different diameters and their pull-times were compared in a balanced design. In addition, because the variation in the diameter of the string can also provide insight into pulling strategy, a variety of end point, topographic and kinematic measures were used to nose position hand pulling movement. These included frame-by-frame video inspection of nose trajectory and hand trajectory using manual and markerless tracking with DeepLabCut and custom written MATLAB scripts that tracked the oscillations of the snout and the left and right hands (Blackwell et al, 2018a; Blackell and Wallace, 2020; Inayat et al, 2020; Singh et al, 2019).

## Methods

### Subjects

Four (2 males, and 2 females) adult negative Thy1-GC aMP mice aged between 8-10 months, weighing 19-30 g were used. The mice were bred and raised at the Canadian Centre for Behavioural Neuroscience Vivarium at the University of Lethbridge. Female mice were housed together and the male mice were singly housed. Animal housing rooms were under a 12:12 light/dark cycle with light-on starting at 7:30AM and temperature set to 21-22°C. All experiments were conducted during the light phase of the cycle. Prior to food-restriction, the mice were weighed over three consecutive days to obtain the average weight at the baseline. On the fourth day, the food was removed, and mice were subsequently maintained on a light food deprivation schedule that maintained their weight at 90% of the baseline weight. Each mouse was weighed and fed daily after training/testing. Mice had access to water *ad libitum*. Behavioral protocols were approved by the University of Lethbridge Animal Welfare Committee and in accordance with guidelines from the Canadian Council of Animal Care.

### Apparatus and training

To familiarize the mice with the strings, short pieces of all types of string used in the experiment were dangled into the mouse’s home cages, and the mice spontaneously pulled them in to incorporated them into their nests. Then the mice were placed in a pretraining apparatus that was a transparent plexiglass tub (46 cm × 26 cm x 26 cm) with a wire mesh top. Stings were dangled into the apparatus so that the mice could pull them in and find the food attached to the end. Figure 1 shows the string-pulling filming box. It was a transparent rectangular plexiglass box (20 cm high x 9 cm wide x 20 cm deep) with no top. The two side walls of the string-pulling box were covered with regular white paper to reduce reflection for filming, and the white paper was placed on the cage floor and was changed each day for each mouse. In the top-middle of the front wall of the box, a ring was attached to maintain the position of the string in the middle of the box during the experiments. One end of the string was passed through this ring into the cage, while the other end was outside the cage with the cheerio attached (Figure 1B). The cage was placed on a table 90 cm above the room floor.

**Figure 1.**
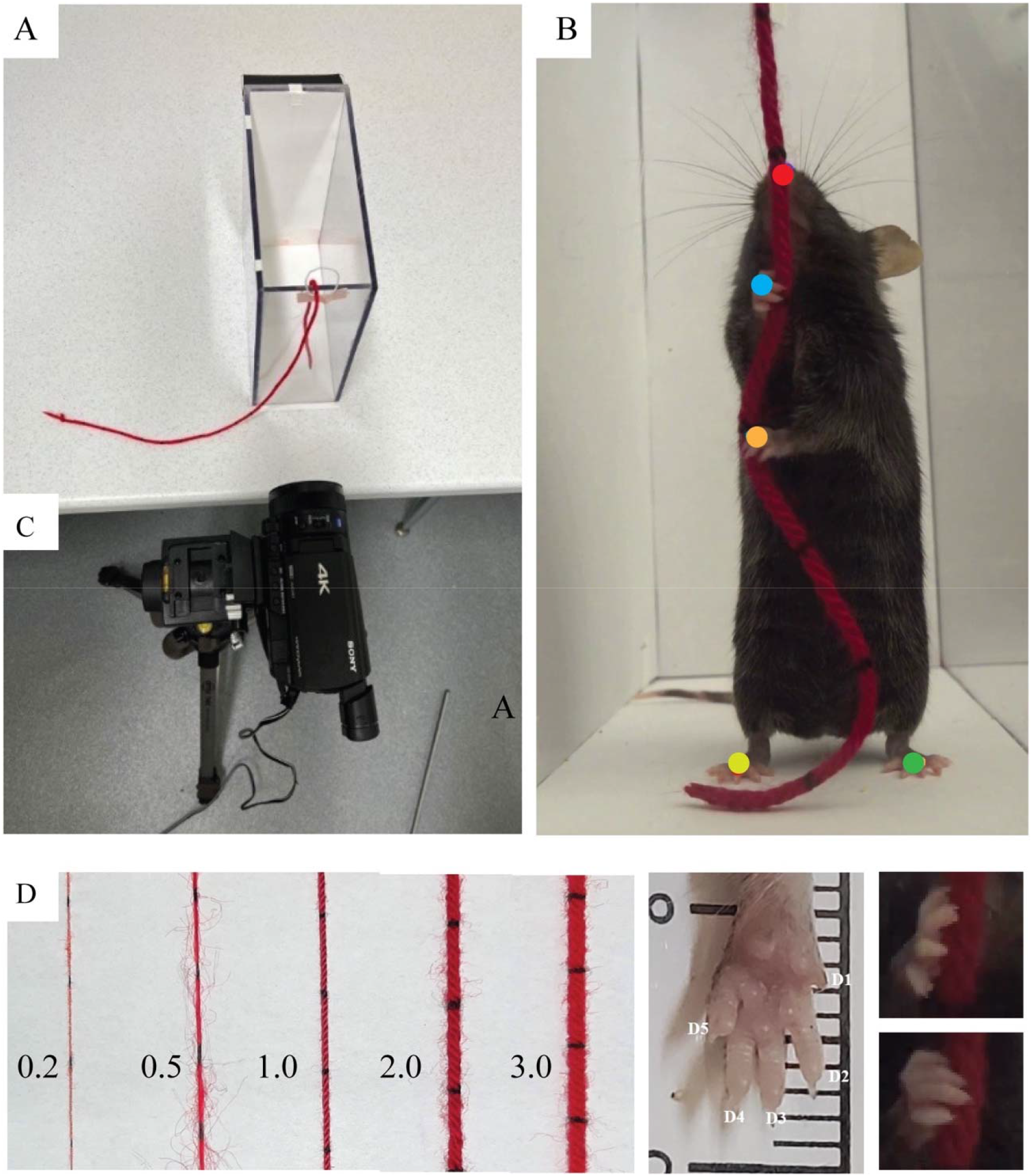
Testing and filming procedure for string-pulling. A. Top view of the test box with the sting inserted 8 cm into to the box. B. Example of a mouse in a standing posture for string-pulling with markers locations for nose and hand tracking. C. Camera location for filming. D. Relative diameters of the five strings that were given to the mice to pull.

### Video recording

Behaviour was filmed using a Sony FDR-AX700 4K HDR Camcorder positioned a fixed location from the test box and facing the front wall of the test box (Figure 1C). The camera settings were adjusted to indoor conditions with a frame rate of 120 frames per second, resolution of 1080 x 1920 pixels, and shutter speed of 1/3000 ms. The recorded videos were transferred to an SD card and later saved on a computer for analysis.

### Strings

Five strings of varying thickness were used (Figure 1D). Each string was 60 cm in length, red in color, and had black marks at 2 cm intervals. String thicknesses were 0.2, 0.5 1.0, 3.0, and 5.0 mm. To start each test, 8 cm of each string was suspended inside the cage in such a way that the mice could rear up and grasp it, while the remaining length of the string was outside the cage. Strings are described as String 1-5, on the basis of increasing size.

## Procedure

First, mice were habituated to different string thickness each day for 10 days in the pretraining box. Each mouse was placed in the holding tub with number of different strings of different diameter (20-30 cm) hanging from the wired mesh top. All mice quickly learned to pull in the strings and consume the food on the end of the strings. Thereafter, the mice were placed in the string-pulling apparatus for five days and given three strings per day to pull, each of which was a different diameter. Pretraining ended when each mouse quickly pulled in the strings. Filming began on the sixteenth day. Each mouse was randomly assigned each string diameter in a semirandom order and testing continued until all mice had pulled in each of the five strings on four trials.

## Video Analysis

The video was first scored manually (Blackwell et al, 2018a). For manual scoring, the videos were played frame-by-frame in Window Media Player (https://support.microsoft.com/en-us/windows/get-windows-media-player-81718e0d-cfce-25b1-aee3-94596b658287) or by Quick time (https://support.apple.com/downloads/quicktime) on an apple computer. Measures of performance included:

*Time*. Time to complete string pulling bout. Time was measured from the first hand contact with the string to the point that the food item struck the floor of the cage as string pulling was completed.

*Reach cycle*. Counts were made of reach cycles with frame-by-frame inspection. A reach cycle was a complete motion by a hand in which the hand was lifted and advanced to grasp the string and then followed by a pull, push and release movement to advance the string (see Blackwell et al, 2020 for reach cycle description).

*Misses*. Misses were counted with frame-by-frame inspection of the video. A miss by either hand was counted when a mouse advanced its hand to grasp a string but failed to do so either by missing the string or by contacting it while failing to make purchase.

*Phase*. The temporal relationship between left-right hand movement and hand-nose coordination was obtained with a wavelet coherence analysis as described by Grinsted et al. (2004). Phase is a measure of the occurrence of the peak frequency of the left hand vs. the right hand and the phase of the hands with the nose, was obtained by comparing the frequency of the left hand, the right hand and the nose.

*Mouth grasps*. Mouth grasps consisting of opening the mouth and grasping the string in an attempt to advance it and were counted using frame-by-frame inspection.

*Kinematic analyses.* The topography of mouth and hand movements including velocity and amplitude of hand movements was documented during each bout with the video analysis tools. The video was analyzed with two computer-based programs, DeepLabCut v2.3 (https://hpc.nih.gov/apps/DeepLabCut.html; Mathis et al, 2018) and MATLAB R2022a (Inayat et al, 2020). The DeepLabCut (DLC) network was trained to track five body parts with 99% accuracy. Body parts include, nose, right and left hands, and right and left feet but only the data from the nose and hands is present here (Figure 1B). The movement of the feet are not reported other than to confirm that the mice largely stood still in the same location relative to the string when string pulling. X– and Y-coordinates of the body parts generated by DLC were used to reconstruct the topographic movement of the nose for each mouse for all string thicknesses. Spatial probability density estimate (nonparametric estimation) of nose movement was generated using kernel smoothing function *ksdensity* in MATLAB. The results of these probability density estimates are plotted as contour plots and normalized 3D spatial probability. Individual reach cycles (lift-advance-grasp-pull-push-release) were identified using approaches described in Singh et. al, 2019 and Inayat et. al. 2020. Briefly, we used Hilbert transform of the y-movements of left/ right hand to identify individual cycles, these cycles were further aligned to the peak y-amplitude, which is the grasp phase. Further, y-coordinates were used to calculate the correlation as a proxy of co-ordination between left and right hand. We then used x/y hand coordinates to calculate velocity profile for each string of varying diameter.

### Statistical Analysis

The statistical analysis of the data was performed using the software IBM SPSS Statistics (v28.0.1.1). The data is also presented as mean ± standard error of mean (Mean ± SEM). For statistical comparisons, a repeated measure ANOVA was used for within-subjects measures (e.g., time, reach cycles, errors etc). A p-value of less <0.05 was defined as statistically significant. Power was measured by eta-squared (η2), a descriptive measure of the strength of association between independent and dependent variables. Follow-up tests were Newman-Keuls tests with significance level set at <0.05.

*Fitts analysis*. The relationship between total pull time (PT) and string thickness was modeled according to Fitts’s law (Fitts, 1954).

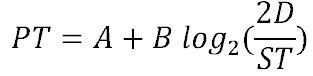

where ST is string thickness, D is the distance from the string, and A and B are model parameters. The log term is usually defined as the “index of difficulty (ID)”. Because the mouse approached all the strings of different thicknesses in a similar manner and stands at a relatively fixed location, the distance from the string was assumed to be constant and the model formulation was modified as follows:

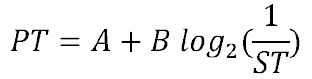

Matlab’s function “fitnlm” for non-linear regression was used for model fitting separately for each mouse as well as for the average PT (over mice). For each mouse, average PT over trials was used. Model appropriateness was tested by fitting the data to linear and quadratic models and comparing F-statistics and p values.

## Results

Because the mice were extensively trained to pull strings before the formal test with the five strings with different diameters, all mice readily and quickly completed reach bouts in which they pulled in a string to obtain the food reward on the end of the string. There were many differences in the way that the mice advanced the strings, as the mice had difficulty with String 1 (0.2 mm dia) (Video 1) and displayed the best performance with String 4 (2.0 mm dia) string (Video 2). The mouse illustrated in Video 1-2 (mouse 1), displayed the best performance of all mice including best performance with the small string. Consequently, time measures distinguished string diameter and other measures including hand cycles and the number of misses and hand shaping, mouth use, and topographic and kinematic changes distinguished motor strategy.

*Hand cycles*. A hand cycle consisted of the movements of lift, advance, grasp, pull, push, and release (Blackwell et al, 2018). The mice made a comparable number of cycles with their left and right hands (right = 1098 vs left 1131) and produced a total of 2,229 cycles in pulling all string diameters in the tests. There was an effect of string diameter on the number of hand cycles, Cycles F(4,12)=7.12, p<0.004, η2=0.84, with more cycles occurring for the smaller than the larger string diameters (String 1 = 17.4±1.7; String 2 = 14.75±0.84; String 3 = 12.9±0.5; String 4 = 12.5±0.7; String 5 = 12.6±0.3; with follow-up tests showing that String 1 and String 2 taking significantly more hand cycles to pull than Strings 3-5.

*Misses*. Misses for the left and right hands are shown in Figure 2A-B. There were significant differences in the number of misses, failure to grasp the string on making a reach as indicated by an effect of String diameter F(4,12)=11.65, p<0.001, η2 = 0.8 (String 1 = 3.28±0.7; String 2 = 1.28±0.21; String 3 = 0.53±0.15; String 4 = 0.15±0.06; String 5 = 0.78±0.27: with follow up tests indicated that String 1 and 2 were associated with significantly more misses than String 4). There was no effect in the number of misses as a function hands, Hands F(2,12)=0.22, p=0.68, η2 = 0.69. There was also no interaction in errors between hands and string diameter, Hands by Strings F(4,36)=0.91, p=0.48, η2=0.24.

**Figure 2.**
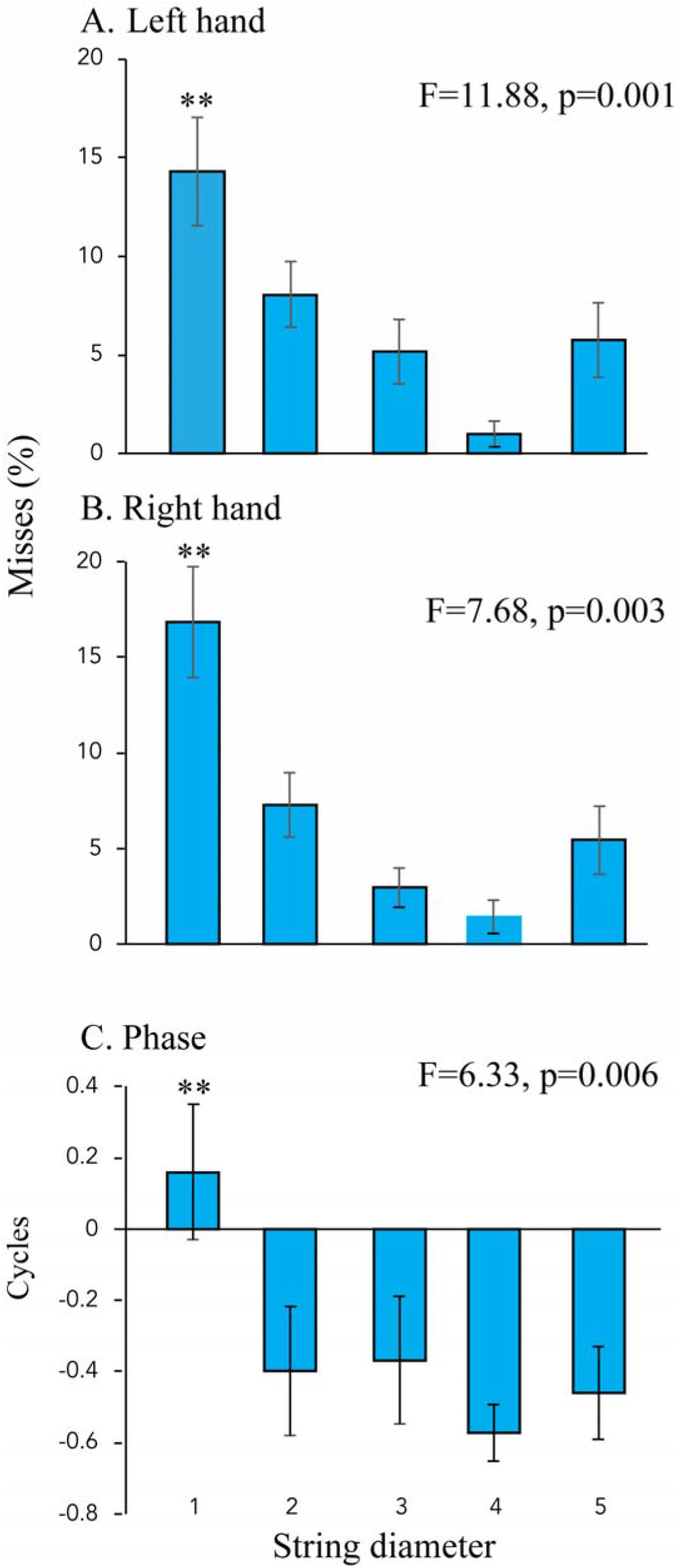
A and B. Error rate for string pulling as measured by misses of the string when grasping by the left hand and the right hand. C. Phase relations between the left and right hands when string pulling. Note: that for the small string the phase is similar as if the mice were pulling with one hand whereas for other stings pulling was performed by out of phase hand over hand movements.

*Misses per hand cycle*. Because misses were defined as failed attempts to pull the string after a reach attempt was made, misses were counted separately for each hand and quantified as a percentage of the total number of attempts using the following formula: Percentage of misses = 100 * misses/(misses + reaches). Figure 2 shows the normalized miss performance as a function of the total number of attempts. A repeated measures ANOVA was used to study the effect of string thickness and trials on the percent of misses. There was an effect of string thickness for both hands (Left Hand: [F(4,12) = 11.88, p < 0.001, η^2^ = .48], Right Hand: [F(4,12) = 7.68, p = .003, η^2^ = .42]) but no effect of trials or any interaction. Posthoc comparisons showed that for left hand, the percentage of misses for String 1 was significantly larger compared to that for String 3-5. Although, there were no significant posthoc comparisons for the right hand, the trend of average values was similar to that for left hand.

*Phase.* Figure 2C shows the phase relationship between the left hand and the right hand were different in relation to string size, F(4,12)=6.33, p=0.006. Follow up tests (p<0.05) showed that String 1 was significantly different from the other string sizes with almost zero phase, as if the mice were pulling the sting with both hands concurrently, which they often did. When the mice pulled the larger strings, the left and right hand were out of phase, indicating that the mice were making alternate pulls with the left and the right hand.

A comparison of the phase relations between the hands and between the hands and the nose are illustrated in Figure 3 for String 1 (Figure 3Ai) and String 4 (Figure 3Bii). We found that for thin string the left– and right-hands co-vary with in-phase movements in the y-direction (up and down movement) as indicated by rightward arrows in Figure 3Aii. Further evaluating the left-right motion (x-direction) of hand and nose reveal that hand-nose also co-varied with in-phase movements as shown in (Figure 3iii). The frequency of these movements were approximately 1 Hz, suggesting that they are both produced by movement of the body. For String 4, the left-right hand co-vary with 180 degrees out-of-phase (anti-correlated) movements as shown by left arrows in Figure 3Bii, and the hand-nose co-vary with in-phase movements as shown by right arrows in Figure 3Biii. These results suggest that hand advance the string with hand-over-hand-movements at about 4Hs and the hands follow the nose movements in x-direction at about 4Hz.

**Figure 3.**
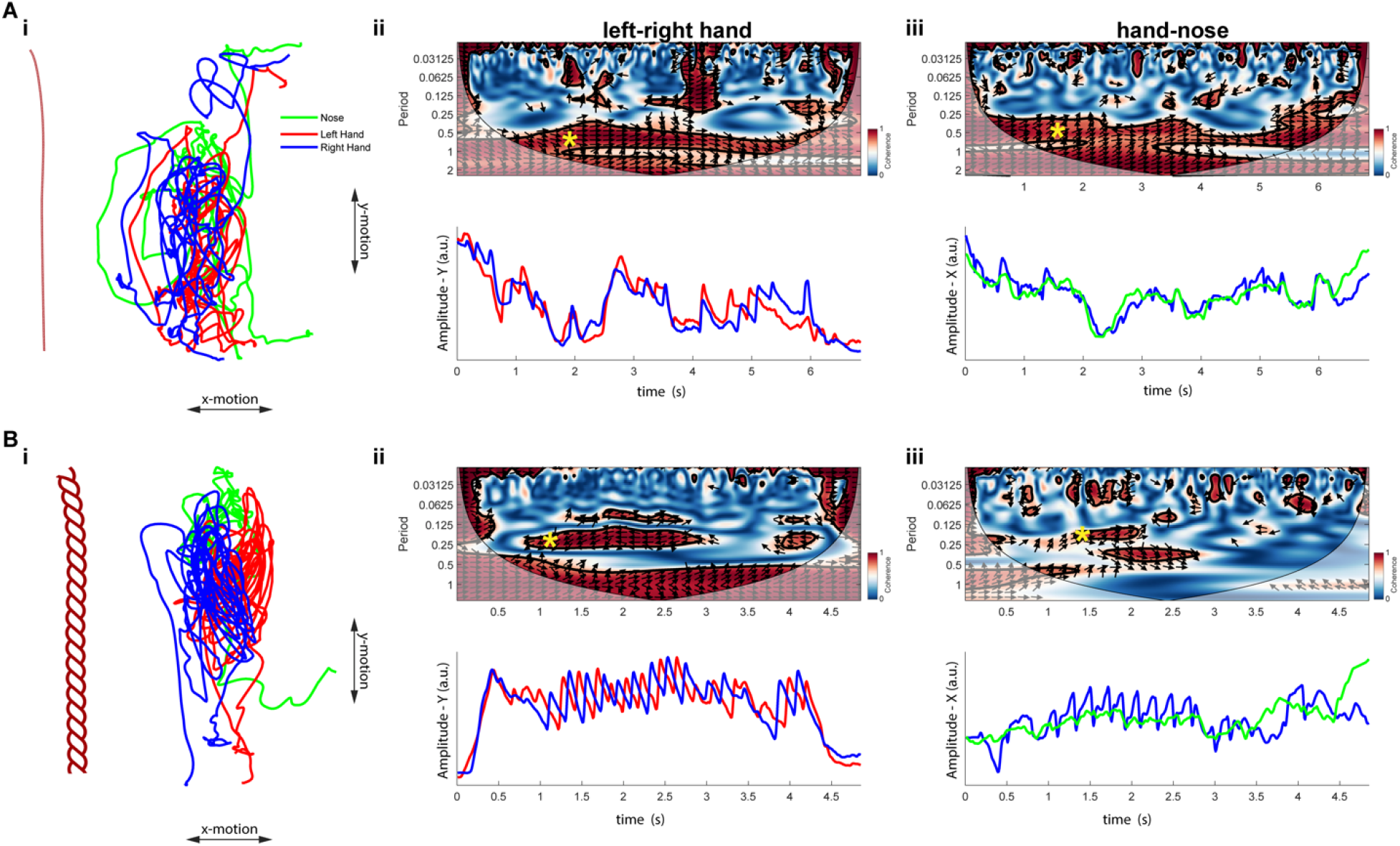
Wavelet coherence analysis of left-right hand coordination and hand-nose coordination for (A) the thinnest string and (B) and the fourth thickest string. (Ai-iii) and (Bi-iii) presents left hand (red), right hand (blue) and nose (green) movements; left-right hand and hand-nose wavelet coherence for thin and thick string respectively. The yellow asterisk on the wavelet spectrogram indicates regions in time frequency space where the two movements co-vary significantly. The significant wavelet coherence levels were determined using Monte Carlo methods. Phase arrows indicate the relative phase relationship between movements (right: in-phase; left: anti-phase; up/down: lead/lag by 90 degrees). A. Note that in phase movement of the hands and the hands and nose have a low frequency of about 1 Hz suggestive of whole body movement. B, Note that the out of phase hand movements at about 4 Hz reflect alternating hand movements whereas the in phase nose and hand movement at 4 Hz suggests that the hand is following the nose.

*Hand shaping*. There were difference in the hand shaping movements the mice used to grasp strings of different diameters. When grasping small strings, the mice could achieve whole hand grasps in which fingers 2-5 pressed the string against the palm, but the string often slipped between two fingers and so was held with fewer fingers. For larger strings, the mice used whole hand grasps, in which fingers 3-4 were proximate and fingers 2 and 5 opened to grasp so that the string was grasp and pressed to the palm with three-point contact. This grasp pattern was assisted with the nails that could be embedded into the string to provide a solid purchase. The following sections describe addition behavioral differences related to string diameter along with topographic representations of movement features. As the following sections describe, the mice took longer to pull thin strings, had trouble tracking the small string with their nose and made many mouth grasps to assist in advancing the string amongst other impairments.

### Pull time, mouth grasps and nose distance

Figure 4 illustrates the trajectory of the nose in a representative mouse pulling each of the five different strings. When pulling down the larger strings, the mouse stands in one location and tracks the string location with its perioral vibrissae as the string moves to the left and then to the right with each hand pull. The heat maps (center) and peak distribution (right) of the nose trajectory indicate that the nose location is more centered and moves less for the thicker strings than for the thinner strings. The same measures show that for thinner strings there is much more nose movement. The increased nose movement is related to the difficulty that the mice have in tracking and in grasping the string and due to their attempts to assist grasping the string with the mouth.

**Figure 4.**
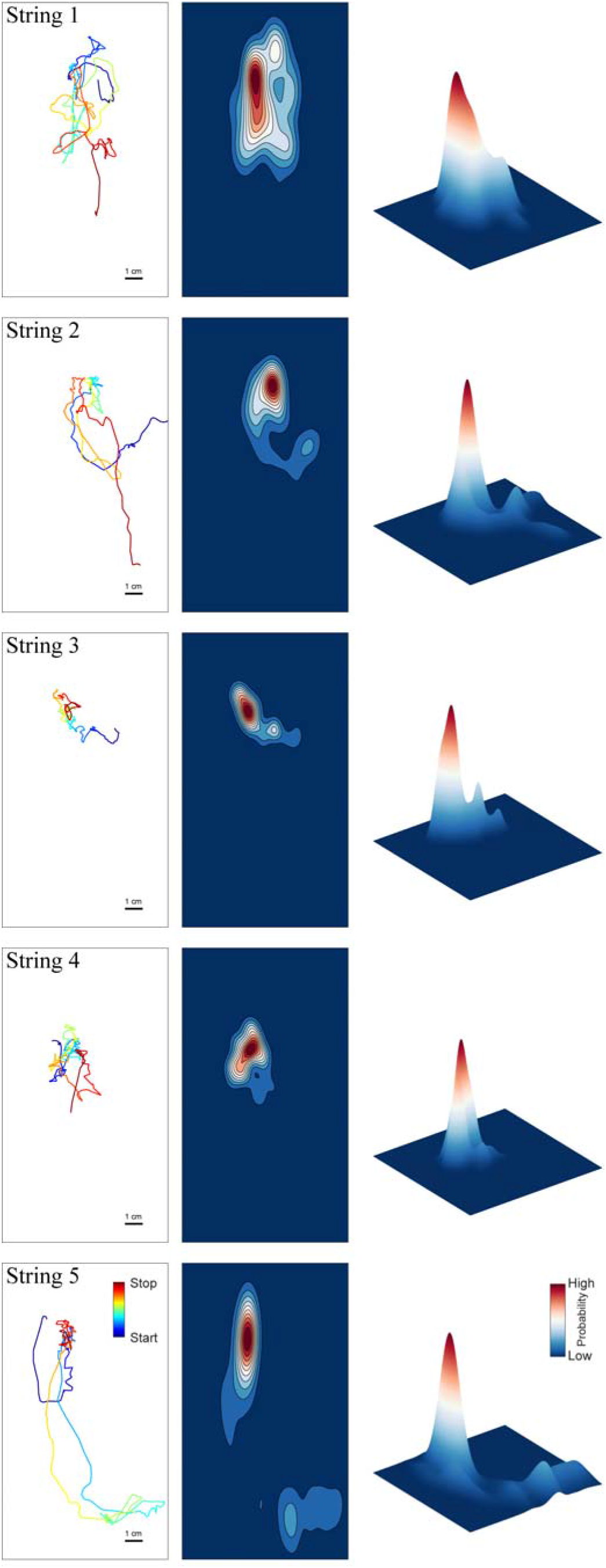
Topographic representations of the nose movements of string-pulling. Left. Movement of the nose color coded for 8 equal division of string-pulling between the start and end of a string-pulling bout. Middle. Contour map of nose movement over the duration of a representative string-pulling cycle for five string diameters. Right. Slope map showing relative nose movement in space.

Figure 5A shows that counts of frame number converted to seconds (120 f/sec) between first contact with the string and the point that the food item attached on its end landed on the cage floor. There was a relationship between string diameter and pull time with the thinner strings taking longer to pull than the thicker strings. This result was confirmed by the ANOVA that gave an effect of Pull time, F(4,12)=15.5, p< 0.001, η2= 0.84. Post-hoc comparisons indicated that String 1 and String 2 took longer to pull than String 4 and 5.

**Figure 5.**
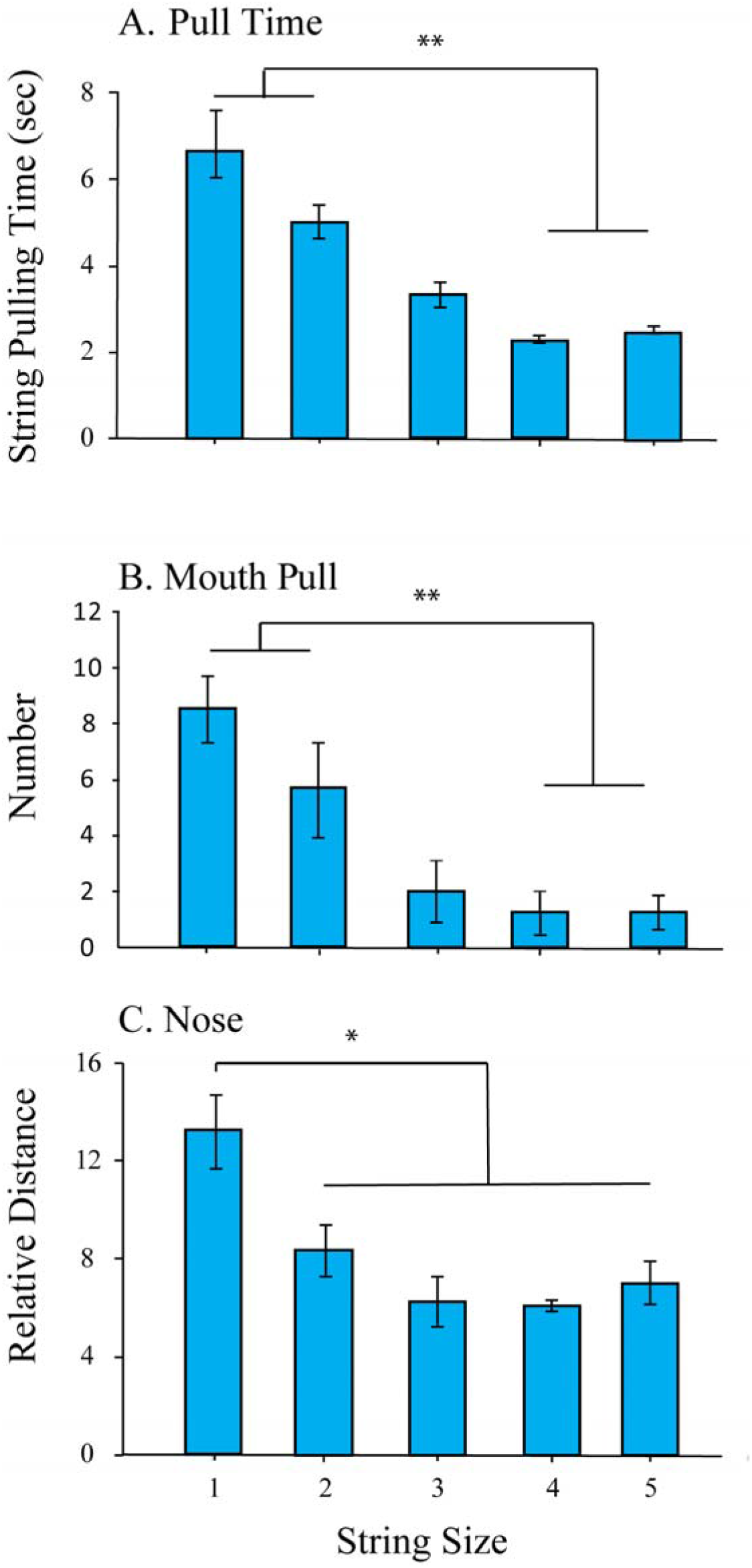
Bar graphs (mean and standard error) associated with strings of five different diameters (1-5 according to increasing diameter). A. pull time. B. mouth pull number. C. Nose movement distance. Note the poorer performance on all measures related to strings of smaller diameter.

Figure 5B shows the counts of the number of times that the mice attempted to advance the string by grasping it by mouth. There was an effect of string diameter and mouth grasps, with the thin strings being grasped more often by mouth than the thick strings, Mouth grasps F(4,12)=23.3, p < 0.001, η2 =0.88. Post-hoc comparisons indicted that String 1 and String 2 were associated with more mouth pulls than String 4 and String 5.

Figure 5C shows the distance travelled by the nose during string-pulling. There was a significant relationship between string diameter and nose movement with greater movement of the nose in tracking the thin strings than for tracking the thick strings, Nose distance F(4,12) = 11.191, p < 0.001, η2 = 0.79. Post-hoc pairwise comparisons indicate that nose travel distance was greater for String 1 than for the other strings.

### Hand amplitude, velocity and fractal analysis

Figure 6 illustrates a MATLAB generated topography of the movement of both hands for complete string-pull sequences from a representative mouse pulling 5 different diameter strings. Each pull features movements of Advance to Grasp the string, Pull, during which the hand holding the strings pulls down to the body midline, Push, in which the hand holding the string pushes the string to the contralateral side of the lower torso, and Release and lift that brings the hand that has released the string back to the body midline in preparation for the next advance (Blackwell et al, 2018). Of these movements, the lift has the highest velocity as indicated by the red portion of the hand trajectory in Figure 4top. The amplitude of the sting pull cycles are similar for both hands and, as is shown in Figure 5-middle, are larger for the thicker strings than for the thinner string. Figure 5-bottom shows hand velocity of the left hand (black curves) and the overall velocity of a complete pull (color curves) are lower for the thinner strings.

**Figure 6.**
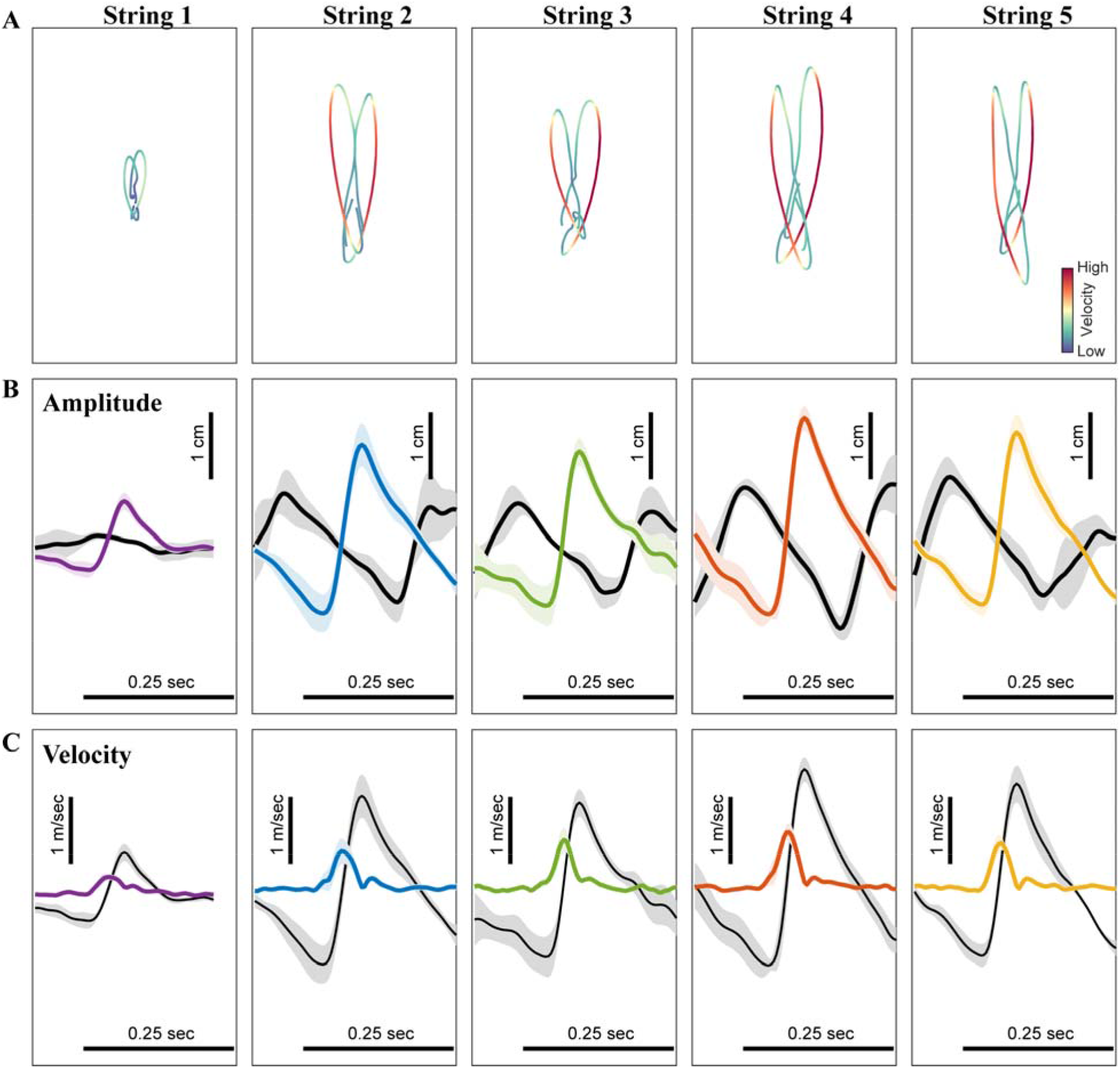
Topographic representations of hand movement for strings of increasing diameter (1-5). The highest velocity indicated by red is the movement of lift with which a hand is raised, after releasing the string, to make a new grasp. A. Topographic representation of hand movement location and speed for strings of increasing diameter (1-5). B. Average hand amplitude for hand movements associated with pulling strings of increasing diameter (1-5). C. Average hand velocity associated with pulling strings of increasing diameter (1-5). Black curves represent mean and standard deviation of upward “lift” movement of the left hand. The color lines give the mean and standard error values for the velocity of all left hand string-pulling cycles for one string-pulling bout.

Figure 7A shows hand amplitude (mean±se) on each cycle of string-pulling as measure by the highest point of grasping to the lowest point of the release. Measures of left hand amplitude gave an effect, Amplitude F(4,12) = 4.120, p = 0.025, η2 = 0.579. Post hoc pairwise comparisons showed that the hand amplitude for String 1 was smaller than was the amplitude for the thicker strings.

**Figure 7.**
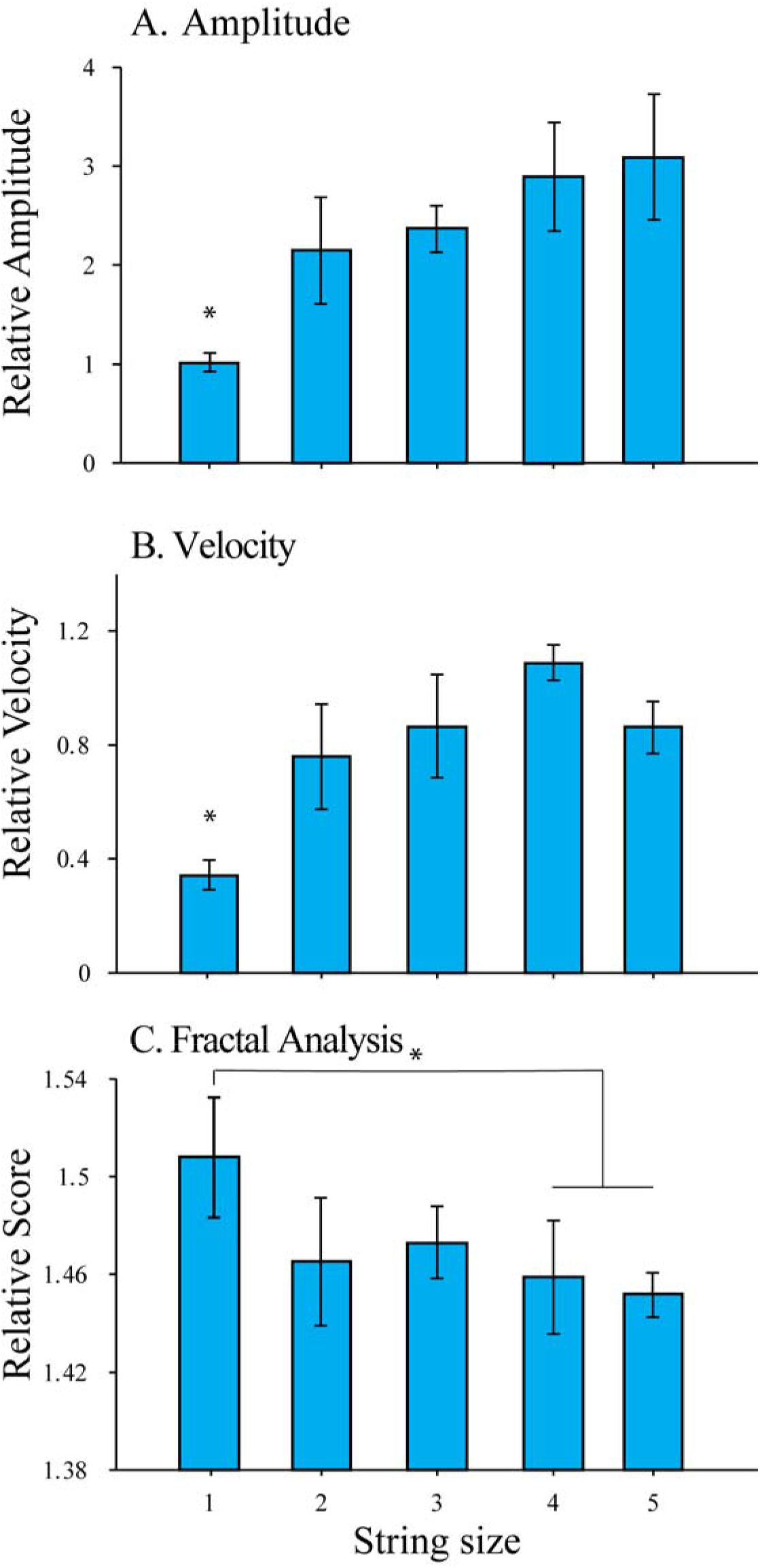
Bar graphs of amplitude, speed and fractal analysis (mean and standard error) associated with pulling strings of five different diameters (1-5). A. Relative amplitude. B. Relative speed. C. Relative fractal score.

Figure 7B shows the velocity of hand movement during string-pulling. The peak velocity in completing a pull cycle is affected by string diameter, Velocity F(4,12) = 6.478, p = 0.005, η2 =0.683. Post-hoc pairwise comparison showed a cycle is significantly slower for String 1 than for the other strings.

Figure 7C summarizes the trajectories travelled by the hands in a pull cycle as measure by a fractal measure. The Hausdorff fractal dimensions were calculated for the nose trajectory using the box-counting method (Inayat et. al., 2020, Singh et. al., 2016). Using this approach fractal dimension is defined as the slope of the line calculated between *log(r) and log(N)*.

Where, *N* is the number of boxes that cover the trajectory, at the magnification ***r***. Trajectories were significantly different, Fractal F(4,12) = 4.375, p = 0.021, η2 =0.593. Post-hoc pairwise comparison among five string thicknesses showed that the fractal value for String 1 was greater than for Sting 4 and 5.

### Total pull time as a function of string thickness follows Fitts’s law

Figure 8 shows the total pull time (PT) versus string thickness (ST) as modeled using a modified formulation of Fitts’s law (see methods). Figure 8 shows average PT (over trials) vs ST and the model is consistent with Fitts’s law as firm lines for each individual mouse which were all significant compared to a constant model, F(1,3) = 16.90, 26.20, 28.12, 20.46, corresponding p = 0.026, 0.014, 0.013, 0.020. Model fitting was also significant for the average PT (over mice) vs ST, F(1,3) = 78.2, p = 0.003. When either a linear or quadratic model fitting was used, both were significant only for mouse # 2, Linear: F(1,3) = 17.19, p = 0.026, Quadratic: F(2,2) = 19.21, p = 0.0495. For the average PT vs ST, the linear model was not significant. However, the quadratic model was significant, F(2,2) = 47.76, p = 0.021, but with a smaller value of F-statistic compared to that for Fitts’s law. These results suggest that the relationship between total pull time and the string thickness followed Fitts’s law. With increasing string thickness, the index of difficulty (log term in Fitts’s law – see methods) reduced and hence the total pull time decreased.

**Figure 8.**
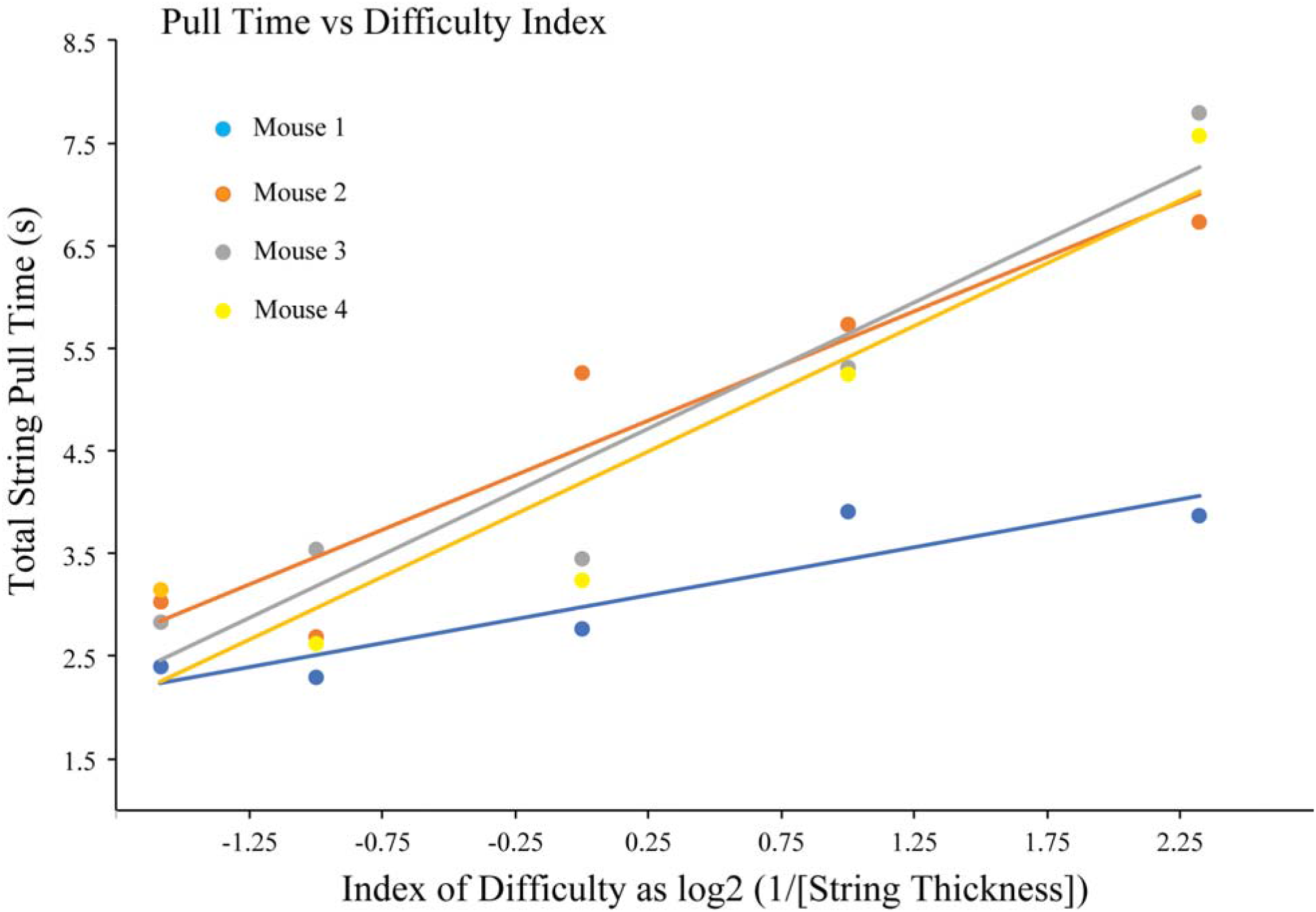
Total pull time as a function of string thickness follows Fitts’s law. Each color corresponds to data from an individual mouse. Isolated markers show average pull time over trials and firm lines show Fitts’s law model fit. Note time is represented as a reciprocal of string thickness.

## Discussion

String-pulling by mice involves a mouse standing on its hind legs and using its hands to grasp and pull in a string that dangles into its holding area. It is a behavior that mice will engage in spontaneously and it is a behavior that can be reinforced with a food item that is attached to the end of the string. The object of the present experiment was to use information theory to assess the time and movements featured in mouse string-pulling by responses to five different diameter strings. End point measures consisted to time to pull and analyses from frame-by-frame video scoring, manual markerless tracking, automatic markerless tracking and video segmentation. Performance in string-pulling was found to vary as a function of string diameter as measured by time, accuracy, nose and hand tracking. The analyses showed that the mice had a proclivity to shift from advancing the string by hand to advancing the string by mouth as string diameter decreased. The temporal results were tested with regression analysis and revealed that Fitts law provided the best fit for performance variations related to string diameter.

### Fitts’s Law

The relationship between movement speed and accuracy, encapsulated by many studies as Fitts’s Law (Fitts, <u>1954</u>), has been used to describe performance in tasks in which a subject makes repetitive movements (Schmidt and Lee, 2020). Such tasks include alternating tapping between two target objects or placing a number of washers on a peg. The Fitts approach has also been used to examine motor cortex function in primates (Iffie et al, 2011). To our knowledge, it has not previously been used to describing mouse motor behavior. The string task lends itself to such an analysis. Maximizing speed and accuracy in making the alternating movements of string pulling maximizes food reward, and so changing task difficulty by manipulating string diameter should be reflected in alternations in time, accuracy and the selection of behavioral strategy. Here we report a significant relationship between string pulling time as a function of string diameter. The application of a number of regression analyses to overall time suggested a best fit that is consistent with Fitts’s law. In the following paragraphs we will discuss some of the strategies and compensatory behaviors that were engaged to cope with task difficulty. Briefly, however, the mice shifted from a hands-only strategy to a mouth-assisted strategy to cope with string diameter change. For the present we suggest a heuristic in subjecting mouse skilled reaching behavior to a Fitts’ analysis is that speed/accuracy tradeoff could be the currency in mastering motor skill for the mouse as it is for humans. In previous studies of mice, rats and humans, we have noted that the relationship between the time of up/down movements diminish across species suggesting that similar approaches could be applied to species comparisons (Singh et al, 2019).

### Mouth use

Ecological specializations are featured in the eating behavior of animals but have been more systematically analyzed in birds than in mammals (<u>Futuyma and Moreno, 1988)</u>. Nevertheless, the analyses of hand *vs* mouth use trade-offs have been recognized in relation to food size. Amongst primates, Peckre et al (2019) report that the strepsirrhines (prosimians) are more likely to use their mouth than their hands to pick up small food items whereas other work (Pouydebat et al, 2008) finds that catarrhines preferentially use a hand to grasp small food items. Here we found that the mice have difficulty in handling the small diameter strings. To cope with the difficulty in pulling smaller diameter strings they revert to using their mouth. Thus, string-pulling presents a paradigm in which there is a time and mouth/hand trade off with respect to target size.

Although it usual for mice to pick up food items with their mouth and then transfer the items to their hands for eating, they do use both their hands and mouth when catching crickets (Glavin et al, 2021). Perhaps this is because crickets are large items and difficult to immobilize, but it is interesting that string variations and movement recreate a somewhat similar hand/mouth trade off problem. It is possible that the diameter of the string itself elicits a mouth grasping response. In studies in which food items of varying diameters are presented to mice, mice are found grab and immediately swallow small items whereas if larger food items are given, they transfer the item to the hands for holding (Whishaw et al, 1990). Perhaps because hand movements and mouth movements are associated as reaching organs with similar neural substrate, their interchangeable use is not surprising (An et al, 2023; Whishaw et al, 2018).

Mouth grasping of the string has also been reported to occur more frequently in rats that display motor abnormalities, suggesting that the mouth is being used as a crutch to assist hand movement (Blackwell et al, 2018). Similarly, anthropoid primates (macaques) revert from hand to mouth pickups after motor cortex damage (Vilensky et al, 1998). Thus, for thin diameter strings, mouth use may be a crutch to deal with difficulty. We have also considered whether mouth grasping could be a result of instinctual drift, in which mouthing the string occurs in anticipation of the food item attached to its end (Breland and Breland, 1996). This possibility seems less likely given that the mice had similar exposure to thin and thick strings, but mainly used their mouth with the thin strings. Whatever the cause of mouth grasping, we favor the view that it is a fallback strategy used by the mouse to cope with task difficulty. Thus, string-pulling and its variations is an interesting test to consider in target populations of the more than 18,500 strains of laboratory mouse (Ju et al, 2022) many of which may show variations in motor system efficiencies (Whishaw et al, 2001).

### Sensory control

For a mouse, as for humans, the string-pulling task is a somatosensory-based task. But mice, unlike humans, orient to the string with their snout, presumably using their perioral vibrissae to locate the string and track its movement changes induced by hand pulls. It is unclear whether sensory information is relayed from the snout to the hands concerning string location, whether the hands learn the location of the string as they pull, and/or whether the mice additionally use the sinus hairs on the ulnar surface of the wrist to locate the string (Carvell and Simons, 2017; Wu et al, 2012). Whatever sensorimotor strategy that the mice do use, they do consistently track the string online with the snout and so the task provides information concerning the perioral sensory abilities of the mouse, information that may be useful for neurological assessment. In addition to providing somatosensory information, the movement of the snout provides information about string-pulling itself. If the snout makes long excursions, it is likely that the mouse has lost the string or has stopped string-pulling and so nose tracking also provides information about the number of pulling cycles that the mice make and the durations of pulling bouts.

When nose movements are tracked with optimal string diameter the nose is associated with a “hot spot” in x-y coordinates that signifies optimal pulling by the mouse. This spot not only signifies optimal pulling, it is also an index of posture. When string pulling, the hands move in diagonal ellipses across the body surface and each excursion must be constrained by posture such that a more upright posture allows a larger excursion. As many neurological and other neural conditions affect posture in addition to skilled movements, nose position in string pulling can give insights into posture *per se* as well as posture related to the execution of a skilled movement task.

### Oscillations

String pulling by mice can be conceived of as the product of two interacting oscillators, one for the nose and/or vibrissae and the other for the hands. As a mouse grasps and pulls a string, the location of the string changes and this change is tracked by movements of the snout. Accordingly, the movement of the snout is in phase with the reaching hand and 180° out of phase with the nonreaching hand. The perioral vibrissae are likely used to track the string’s location but it unclear whether this information is responsible for the movement of the snout only or is conveyed to hands enabling them to locate the string on each reach. It is known that the vibrissae whisking is the product of an oscillator (Takatoh et al, 2022) that produces rhythmical movements of about 7Hz, but the head movement and hand movements have a frequency that is lower, about 4 Hz. Whisking is reported to contribute to the guidance of the forelimbs of mice more generally (Bergmann et al, 2022) and so it is possible that it also contributes to reaching for the string. The rhythmical movements of the hands, which are out of phase are likely also controlled an oscillator (Wagner et al, 2021), but what is not known is whether information from hand contact from the string also signals the head/whisking networks. Clearly, the string pulling task lends itself to the investigation of possible interactions between the neural systems that are instructed by head orientation and whisking and the neural systems that produce skilled hand reaching.

### Methodological considerations

The methodological target of the present study lay in tracking three body parts, the nose, the left hand and the right hand with AI methodology supplemented by visual analyses. Such simplified topographic methodology has been found useful even when tracking a single point in the many thousands of studies of spatial navigation in rodents. This analysis methodology allowed an analysis of tradeoffs between mouth vs. hand use in relation to task difficulty. In addition, the methodology allowed many details of behavior to be simplified into a single comparison, in this case Fitts’s Law. It is important to note that all thicker strings were handled well, but the best string-pulling response was obtained with the 2mm diameter string. Because the availability of string types and diameters is high, the present results suggest that for future studies, consideration of string diameter and type is important. Optimal string thickness and type can be tested by presenting strings to mice in a holding cage and measured by time taken to pull using procedures such as that described here. In addition, as many experimental designs might require variation in task difficulty, string diameter and other modifications of string composition can be used as a task variables.

String-pulling recommends itself as a task for mouse skilled movement assessment because it is a task that mice will spontaneously perform and a task that can be reinforced to obtain reliable performance. Here we found that mice will quickly pull in 60 cm length strings and in doing so would make about 10-15 reach/grasps with each hand. Thus, when give 4 strings to pull each day, they would make about 80 reaches and grasps in each session. This number of grasps is on the high end numerically for tasks in which mice have been trained to reach for food or for water as a method for measuring hand function (Baird et al, 2021; Galiñanes et al, 2018; Klein and Dunnett, 2012; Whishaw and Pellis, 1990; Whishaw, 1996; Wang et al, 2022). The additional advantage of string-pulling is that whereas in most other reaching-based tasks only one hand is trained/assessed, the string-pulling task concurrently requires similar movements from both hands. In the present study, about 40-60 reach/grasps in pulling the four strings were obtained with each hand on each test day from each mouse. In addition, both hands are assessed in a task that requires bilateral hand coordination, an assessment methodology that is difficult to obtain and measure, e.g., as for bilateral food handling while eating (An et al, 2023; Barrett et al, 2022; Whishaw et al, 2018) or bilateral forelimb use when walking on flat surfaces, beams or a rotarod (Carter et al, 2001; Clarke and Still, 1999; Sheppard er al, 2022). A number of reviews have summarized the utility of the study of bilateral motor tasks for the study of motor control and rehabilitation in humans (Darevsky, 2023; Carson, 2005; Waller and Whitall, 2008) and string-pulling can serve a similar experimental function in both mice and humans.

Depending upon filming conditions and the quality of film one or another procedure describe here can be used. The MATLAB video procedure does require good and consistent video records (see Inayat et al, 2020 for a discussion), but the DeepLabCut procedure is sensitive to wider variation in film quality. Thus, depending upon the conditions under which data is collected, the procedure here of tracking the nose and the hands can provide robust results concerning string-pulling. In addition, it should be noted that the simplest measure of performance is time. Time is a limited and valuable resource and animals must use time strategically to ensure survival (Gause, 2019; Huston et al, 1993; Whishaw et al, 1990). Even in string pulling, an error is lost time. Thus, the results of the present study confirm the many studies of spatial navigation that time measures proved a useful summary of performance.

## Conclusion

Here we describe the movements of the nose and hands as a method of assessing string-pulling in the mouse. We propose that because sting pulling speed/duration can be describe by Fitts’s law, the mouse lends itself to studies related to information theory. We also suggest that string-pulling diameter variation can alter task difficulty and assess mouth/hand tradeoffs related to target size. The tracking procedures used here gave very similar information about hand use and trajectory for both optimal and compensatory movements of string-pulling. The effect of string diameter is also described and it provides variation in task difficulty, as the mice had greater difficulty in pulling thin strings than thick strings. String pulling can be used to investigate oscillatory movements of the head/vibrissae and the hands that produce pulling movements. Finally, string-pulling can be useful for the study of information theory as is demonstrated by time fitting to Fitts’s Law. As an aside, the present results are relevant to Pliny’s performance in BF Skinners chain pulling task (see introduction) because Pliny’s use of both teeth and hands suggests that chain pulling was difficult.

## Contributions and acknowledgements

PSS performed the experimental procedures with the supervision of BMA; SA, SS, HRS contributed to the data analysis; MHM provided laboratory facilities and funding; IQW, SA and SS provided the conceptual background for the study, PSS, BMA, SA, SS and IQW contributed to manuscript preparation. This work was supported by the Natural Sciences and Engineering Research Council of Canada (grant# 40352), Alberta Innovates, Alberta Prion Research Institute (grant 43568) and Canadian Institute for Health Research (grant 390930). The authors thank Di Shao for animal breeding.

## Figures

Video 1. A reach bout in which Mouse 1 reels in a 0.2 mm diameter string (the thinnest string) and assists reach cycles by using a variety of compensatory movements and the mouth to pull.

Video 2. A reach bout in which a Mouse 1 reels in a 2.0 mm diameter string (fourth largest diameter string) and mainly advances the string with rapid hand cycles.

